# A biophysical computational model for memory trace transfer from hippocampus to neocortex

**DOI:** 10.1101/503524

**Authors:** Xin Liu, Duygu Kuzum

## Abstract

The hippocampus plays important roles in memory formation and retrieval through sharp-wave-ripples. Recent studies have shown that certain neuron populations in the prefrontal cortex exhibit coordinated reactivations during awake ripple events. Also, the reactivation seems stronger during initial awake learning. These experimental findings suggest that the awake ripple is an important biomarker, through which the hippocampus interacts with the neocortex to assist the memory formation and retrieval. However, the computational mechanisms of this ripple based hippocampal-cortical coordination are still not clear. In this work, we build a biophysical model that includes both CA1 and layer V networks of the prefrontal cortex to investigate the possible mechanisms, by which the memory traces in the hippocampus can be transferred to prefrontal cortex. We first show that the local field potentials generated in the hippocampus and prefrontal cortex exhibit ripple range activities that are consistent with the recent experimental studies. Then, we find that the sequence information stored in the hippocampus can be successfully transferred to the prefrontal cortex recurrent networks through spike-timing dependent plasticity (STDP) and sequence replays. Further, we investigate the mechanisms of memory retrieval in the PFC network. Our findings suggest that the stored memory traces in the prefrontal cortex network can be retrieved through two different mechanisms, namely the cell-specific input and non-specific spontaneous background noise. Finally, we show that more SWRs and an optimal background noise level will both contribute to better sequence reactivations in the PFC network during memory retrieval. Our study presents a possible explanation for the memory trace transfer from the hippocampus to the neocortex through ripple coupling in awake states and reports two different mechanisms by which the stored memory traces can be successfully retrieved.

**Author Summary:** The hippocampal-cortical interactions have been found to be important for learning and memory. Recent experimental work reports interesting coordinations between the hippocampus and neocortex during ripple events in quiet awake states. Investigating this phenomena is critical for a deeper understanding of the mechanisms by which hippocampus contributes to the long-term memory formation in the neocortex. To this end, we build a biophysical computational model for both the hippocampal CA1 and prefrontal cortex networks. We demonstrate that under STDP rule and sequential inputs, the memory traces initially stored in the hippocampus can be transferred to the neocortex through the ripple events. The stored memory traces in the PFC network can be reactivated by two different mechanisms, namely the cell-specific inputs and non-specific background noises. Our study suggests that both the ripple event numbers and the level of background noises jointly determine the quality of memory retrieval during reactivation.

## Introduction

The hippocampus (HPC) plays important roles in memory consolidation and sharp-wave ripples (SWR) are believed to transfer the compressed temporary information stored in the hippocampus to the distributed cortical networks [1, 2] through the abundant connections between the hippocampus and cortex. In support of this idea, it has been shown that during sleep, the neurons in the visual cortex exhibit coordinated reactivation along with the HPC SWR [3] and the neurons in prefrontal cortex (PFC) also display learning-dependent reactivations when SWR are generated during sleep [4]. Furthermore, the firing of PFC neurons falls within the plasticity time window after HPC SWR occurs [5]. Recently, it has also been reported that the neurons in the anterior cingulate cortex show increased correlated activities before HPC SWR [6]. Enhancing the oscillation coupling between HPC and PFC boosts the memory task performance [7]. These experimental findings support the view that HPC indeed play important roles in memory consolidation through SWR during sleep.

Besides sleep, SWR and sequence replay also occur in the HPC during awake immobility [8-10]. Similar with sleep SWR, interruption of awake SWR during spatial learning degrades the animal’s performance in later spatial tasks [11]. Recent studies demonstrate strong hippocampus-cortical modulations during awake SWR. It has been shown that the PFC neurons reactivate during awake SWR and different excitation and inhibition patterns are observed in the PFC neuron populations [12]. Also, the CA1-PFC reactivation is found to be stronger during awake SWR than during sleep SWR. Especially when the animal is learning novel information, the CA1-PFC reactivation is further enhanced [13]. These observations lead to the postulation that HPC not only contributes to memory consolidation during sleep, but also takes active part in the hippocampus-dependent memory retrieval and initial learning process during awake behavior. The mechanisms for the HPC SWRs assist memory formation in the PFC are poorly understood.

Here, we build a biophysical computational model that includes both a CA1 network [14-16] and a PFC network [17] to study the memory transfer from the HPC to the PFC. Under the sequential inputs from a virtual CA3 network, the CA1 network generates ripple range oscillations along with simultaneous sequential pyramidal cell reactivations. The firing activity in CA1 potentiates the pyramidal cells in PFC through the monosynaptic connections to induce coordinated sequence reactivations. By enabling STDP between PFC neurons, we demonstrate that the sequence can be stored in the recurrent connections of the PFC network. Later, we find that the stored memory traces can be reactivated through two different mechanisms, namely the cell-specific local stimulation and non-specific spontaneous background noise. Our modeling results provide a possible explanation for the hippocampus dependent memory formation in the cortex during SWR in awake state.

## Results and Discussion

### CA1 and PFC network activities during memory trace transfer

The CA1/PFC coupling network model is illustrated in the schematic shown in Fig 1. The CA1 network consists of 400 pyramidal (PY) neurons and 100 basket (BS) cells. The PY neurons project to BS cells through AMPA synapses and receive GABA projections from the BS cells. The gap junctions exist between adjacent BS cells. For the PFC network, there are 100 PY neurons and 25 interneurons (IN). The PY neurons form unidirectional and bidirectional AMPA synapses on each other and also send excitatory inputs to the IN neurons. The IN neurons connect back to the PY cells through GABA synapses and also form recurrent inhibitory connections between each other. The details of the network model are explained in the methods section.

**Figure 1.**
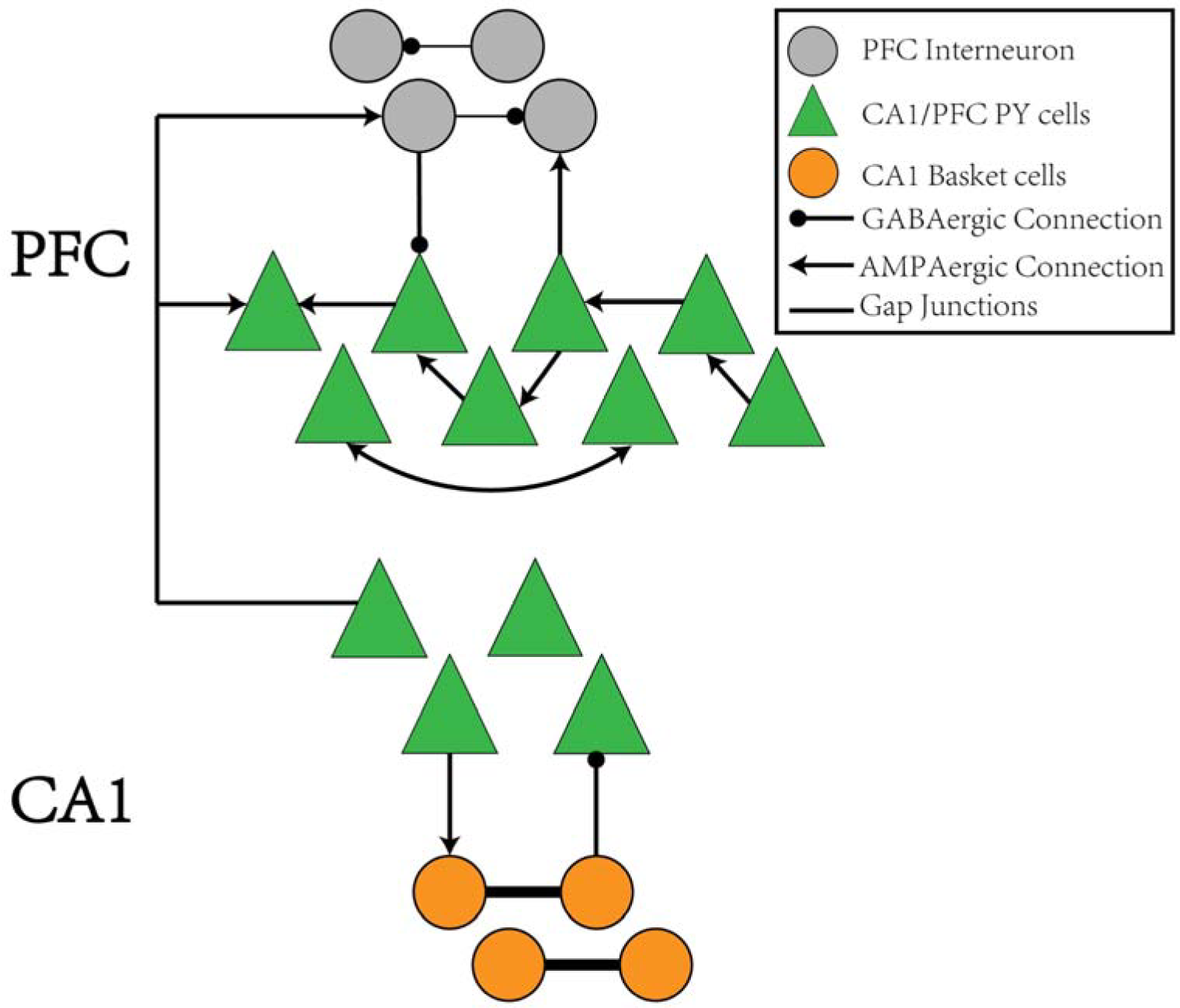
The schematic of the CA1-PFC model. For the CA1 network, the PY cells connects to the IN cells with AMPA synapses, whereas the IN cells project back to the PY cells with GABAa synapses. There is no synaptic connections between the PY cells in CA1. For the PFC network, the PY cells forms unidirectional and bidirectional recurrent connections among each other with a certain probability. The PY cells also project to IN cells with AMPA synapses. The IN cells in PFC have recurrent GABAa connections among each other and project back to the PY cells in PFC through GABAa synapses, Between the CA1 and PFC networks, the PY cells in PFC receive AMPA synaptic inputs from the PY cells in CA1 network. The IN cells in PFC also receive AMPA synaptic inputs from PY cells in CA1.

To mimic the CA3 input that drives the CA1 network, we give sequential noisy inputs (see methods) to both the PY neurons and BS neurons in the CA1 network. In this occasion, the PY cells in the CA1 network get depolarized and fire sequentially, as shown in Figs 2a and 2b. As a consequence, the local field potential (LFP) recordings in the soma layer show ripple transients overlapped on the sharp-waves (Figs 2c and 2d). Fig. 2e is a spectrogram of the LFP recordings in the CA1 network, showing ripple range oscillations around 200 Hz.

**Figure 2.**
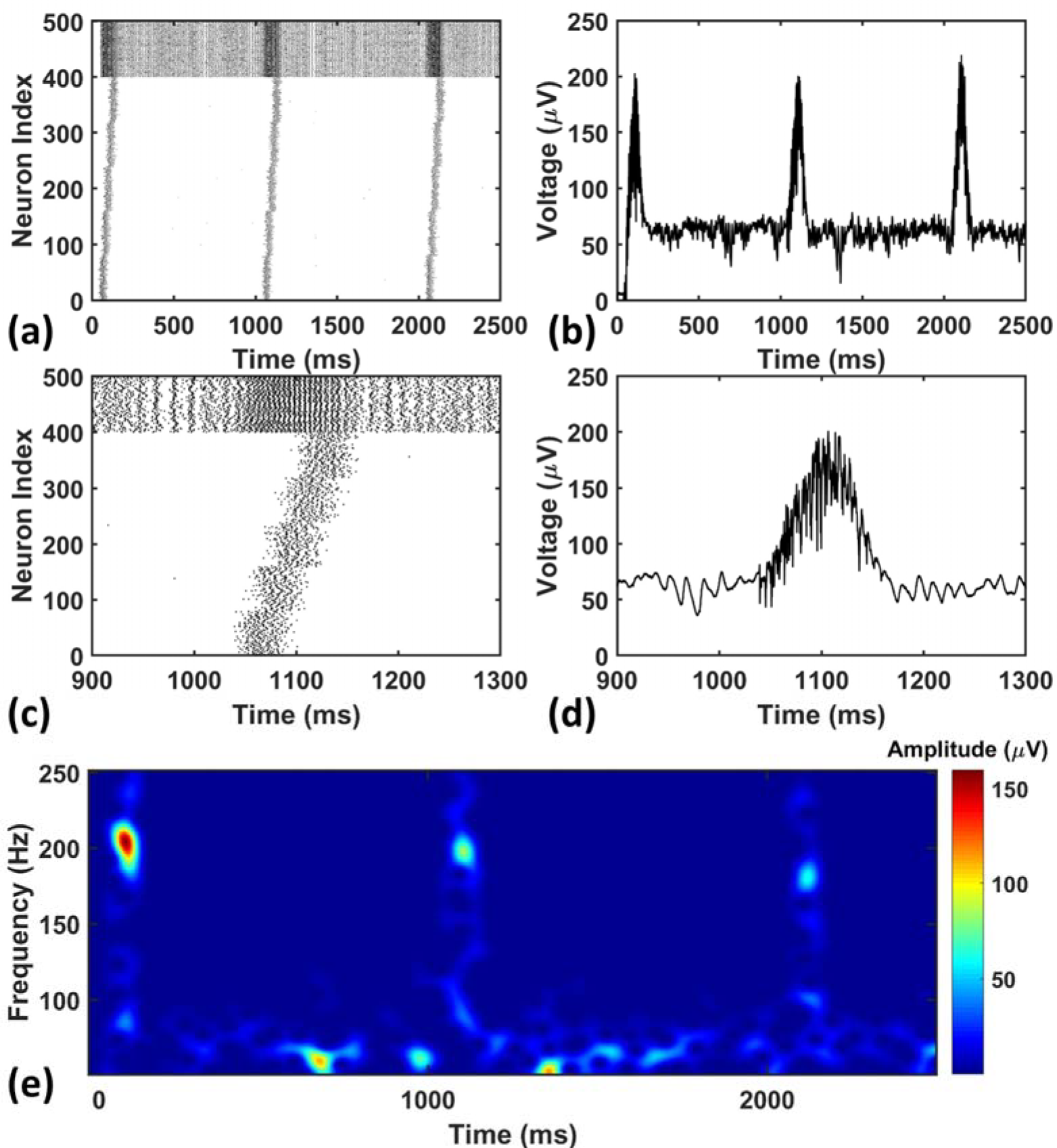
The raster plot and LFP recordings in CA1 network. (a) The raster plot of the CA1 network during the simulation. Under the sequential noisy inputs, the PY cells in CA1 network shows ordered firing across 5 different groups. (b) The simultaneously recorded LFP shows ripples and sharp waves. (c) A zoom-in raster plot of one ripple event in the CA1 network. (d) A zoom-in LFP recording during the ripple event shown in (c). (e) A spectrogram of the LFP recordings showing ripple range oscillations during sequential replay in the CA1 network.

For the PFC network, when the CA1 PY cells fire and send excitatory inputs to both the PY cells and IN cells in PFC through monosynaptic connections, the PY cells in PFC fire sequentially (Figs 3a and 3b). The IN cells also increase their firing frequencies. In the same time, as shown in Figs 3c and 3d, the LFP recordings in the PFC network exhibit transient oscillations that have components in high gamma (60-100 Hz) and ripple range (100-250 Hz). This can also be seen from the spectrogram of the LFP recordings in PFC network (Fig. 3e).

**Figure 3.**
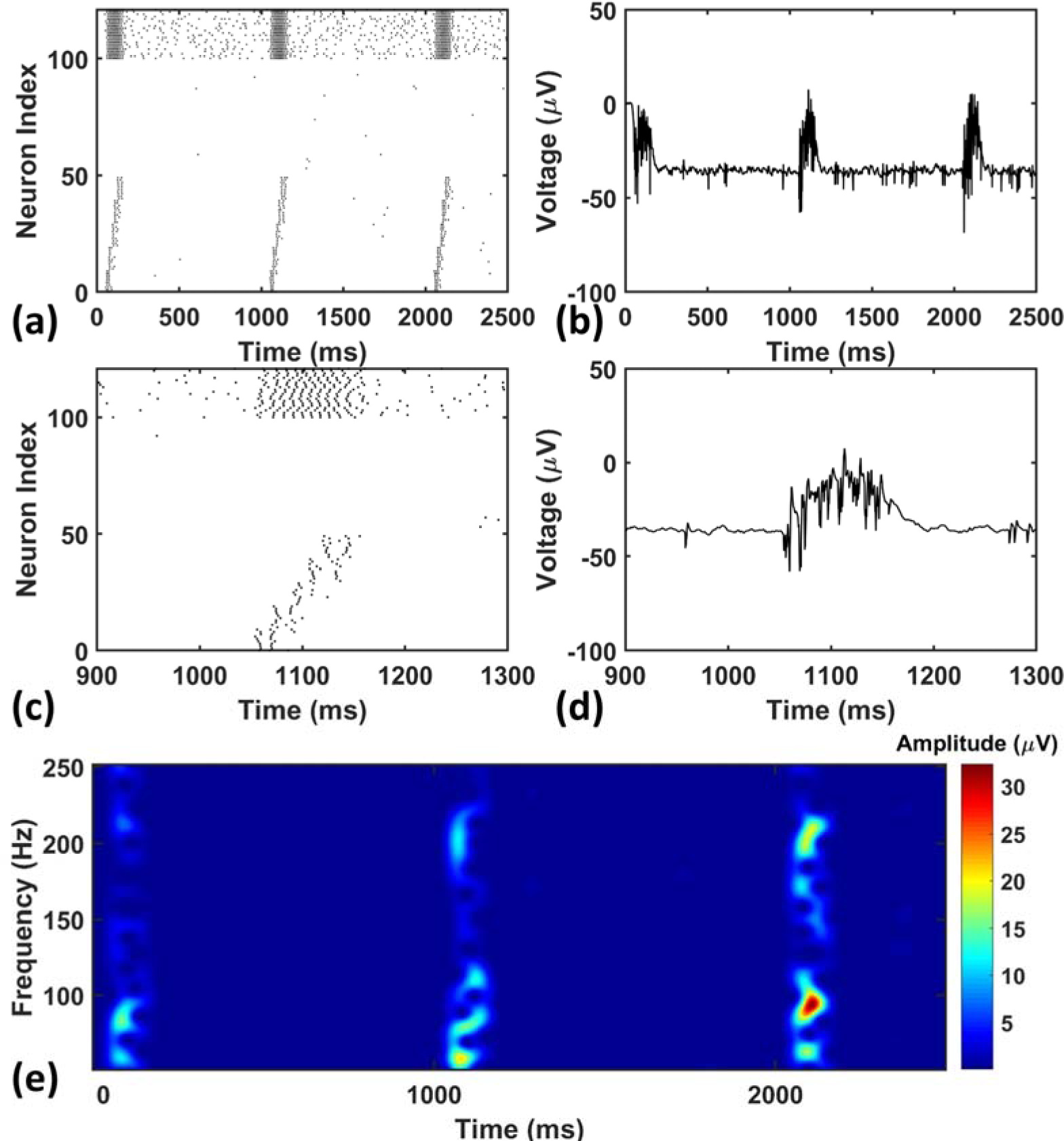
The raster plot and LFP recordings in PFC network. (a) The raster plot of the PFC network during the simulation. Under the monosynaptic inputs from CA1, the PY cells in PFC network shows sequential firing. (b) The simultaneously recorded LFP in the PFC network. (c) A zoom-in raster plot of the PFC network during the ripple event in CA1. (d) A zoom-in LFP recording in the PFC network during the sequential firing shown in (c). (e) A spectrogram of the LFP recordings showing high gamma and ripple range oscillations during the CA1 ripple events.

Recent studies on HPC-PFC interactions have revealed various neural activity modulations in both awake and sleep state [18]. During sleep, it has been shown that the localized ripple oscillations detected in the LFP recordings at PFC are strongly coupled to the ripple events in the hippocampus [19]. Also, the coupling strength gets stronger after the animal performs a spatial learning task. Our LFP simulation results are consistent with these observations, suggesting that the ripple-ripple cross-frequency coupling might serve as a communication link between the cortex and hippocampus for memory trace transfer.

### The sequence transfer in the hippocampus-PFC network

Next, we investigate the possibility of the memory transfer through CA1-PFC communication and STDP. The CA1 network generates 5 ripples per second and the simulation time is set to 3 seconds. The raster plot and LFP recordings in PFC from one representative simulation are shown in Fig 3. When the CA1 network generates ripples, parts of the PY cells in PFC fire sequentially, while all the interneurons increase their firing rate compared with no-ripple time intervals. Due to the STDP rule, the PY-PY connections and IN-PY connections are updated according to the spike timing of the cells (Fig 4a and 4c). As shown in Fig 4b, by the end of one representative simulation (3000 ms training time), the feed-forward AMPA connections among the PY cells that receive direct CA1 inputs are strengthened, whereas the feed-backward AMPA connections are weakened. Since the inhibitory STDP rule is symmetric and LTP-only, all the IN-PY GABAa connections for the PY cells that receive direct CA1 inputs are stronger by the end of the simulation (Fig 4d). However, for the PY cells that do not directly receive CA1 input, the strengths of their GABAa synapses are almost unchanged because of their sparse firing activities. Therefore, under the STDP rule for both excitatory and inhibitory synapses in the PFC network, the sequence initiated in CA1 is successfully transferred to the PFC network and stored in the recurrent synaptic connections of the cortical cell populations.

**Figure 4.**
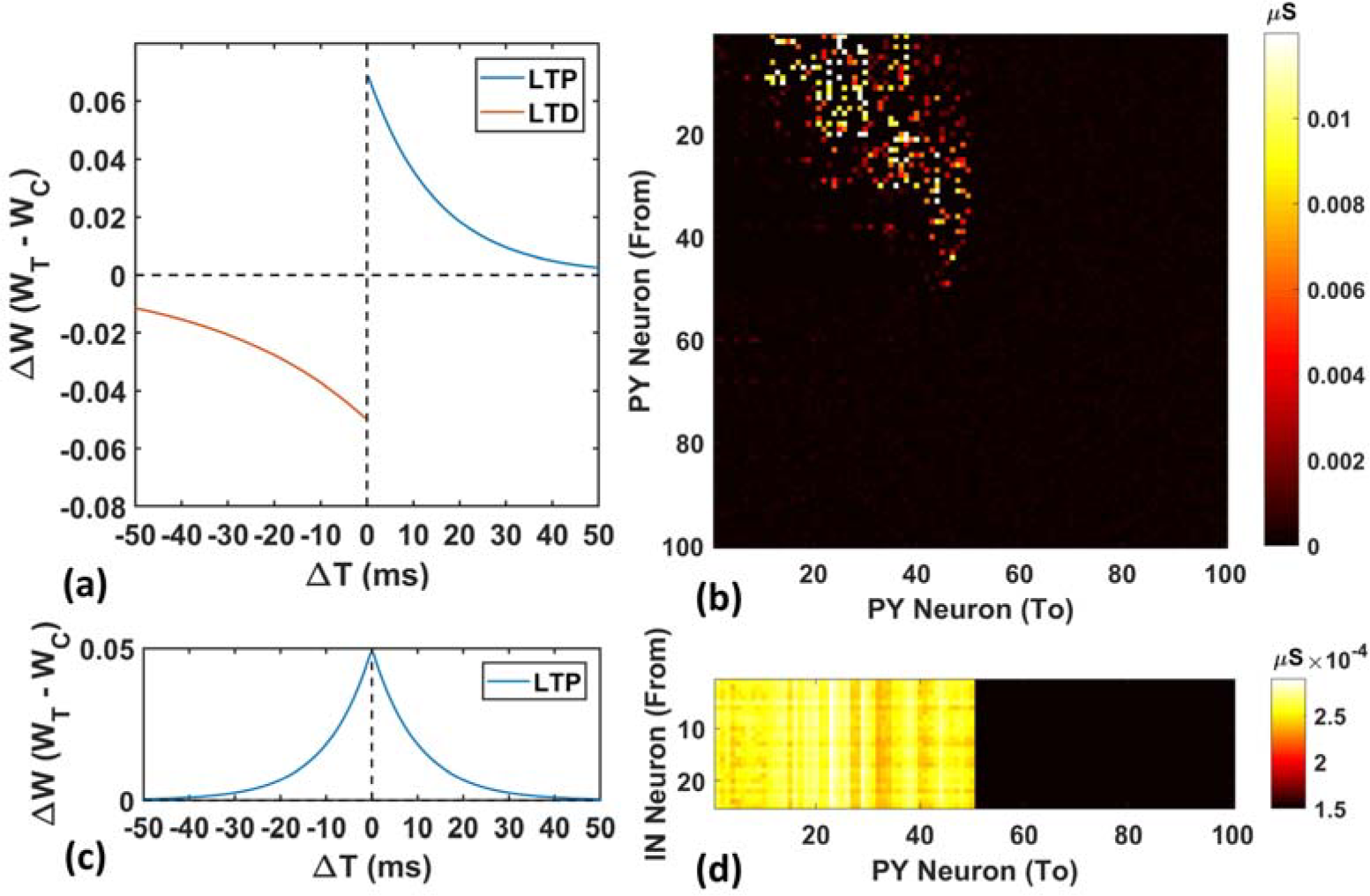
The weight matrices of the PFC network. (a) The STDP weight update rule for the PY-PY AMPA synapses in the PFC network. (b) The weight matrix of the PY-PY AMPA connections after learning. It can be seen that, for the PY cells that receive CA1 input, the feed-forward connections among them get strengthened, while the feed-backward connections remains weak. (c) The STDP weight update rule for the IN-PY GABAa synapses in the PFC network. (d) The weight matrix of the IN-PY GABAa connections after learning. Since the first 50 PY cells in PFC network fire sequentially during the learning, all of the corresponding IN-PY connections are strengthened due to the symmetric LTP-only STDP rule for GABAa synapses, whereas the rest connections does not get potentiated much.

### The sequence replay in PFC network induced by cell-specific input

After the successful storage of the CA1 sequence, we investigate the retrieval of the memory in the PFC network by cell-specific stimulations to part of the PY cells in PFC network. The cell-specific input induced replay represents the scenarios where the memory is recalled by the sensory stimulation or natural cues, which is commonly observed in studies on contextual fear conditioning [20].

To apply the cell-specific input, we inject a train of 5 Hz 5ms duration 0.7 nA step currents into the soma of the first 10 PY cells in the trained PFC network and compute the Matching Index (MI) of sequence replay (noise level = 0.053 nA^2^, see the Methods). The MI value quantitatively measures the replay quality. A perfect forward replay will have a MI value of 1, whereas the random replay will have a MI value of 0. We also perform simulations in the untrained PFC network using the same stimulation protocol as control group. The raster plot of the trained PFC network is shown in Fig. 5a. It can be seen that, under the current injection, the first 10 PY cells fire and quickly recruit the rest of the downstream PY cells to form a sequence in a temporally compressed manner (~30 ms). This replay mechanism relies on the strengthened AMPA synapses between PY cells and a sufficient number of recurrent PY-PY AMPA connections. The firing of each PY cell will elicit an excitatory post-synaptic potentials (EPSPs) in the downstream PY cells, which elevates the membrane potential. If the postsynaptic PY cell receives multiple EPSPs from multiple presynaptic PY cells in a short time interval, it is more likely to fire. For the untrained network case, the raster plot is shown in Fig. 5b. It can be seen that even though the first 10 PY cells fire under the external stimulation, the rest of the PY cells do not fire to form a sequence. This is due to the weak PY-PY AMPA connections in the untrained network such that the firing of first 10 PY cells cannot generate large enough EPSPs in the downstream PY cells. To further quantitatively explore the possible factors that can affect the sequence replay, we examine two parameters that can change the sequence MI in PFC network: the AMPA noisy input level and the recurrent AMPA connection degradation.

**Figure 5.**
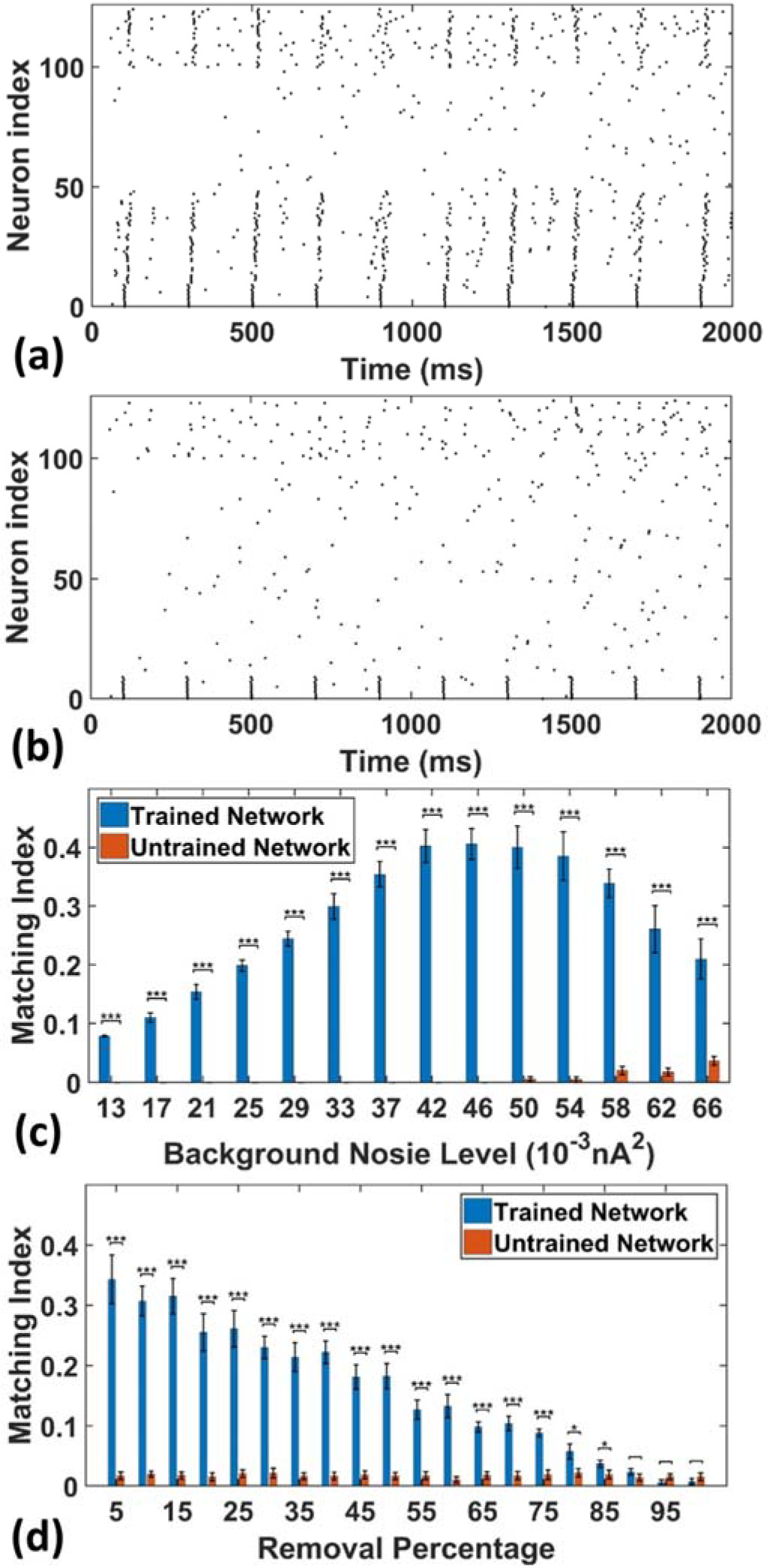
The sequence replay in PFC network induced by cell-specific current injection. (a) When the first 10 PY cells in the trained PFC network are given a short 5 ms current injection, the downstream PY cells are recruited by the feed-forward AMPA projections and a sequence is induced (noise level = 0.053 nA^2^). (b) For the untrained PFC network, the PY cells exhibit random firing and the activation of first 10 PY cells do not induce sequence replay. (c) The sequence matching index MI induced by strong cell-specific input depends on background noisy input to the PY cells. Note that the MIs of sequence replay in trained network are significantly different from those in the untrained network for all the noise input levels (rank sum test). (d) The sequence replay MIs depend on the PY-PY connection integrity. Massive loss of AMPA connections between PY cells will result in insufficient EPSPs to depotentiate the membrane potentials of downstream PY cells, which will in turn suppress the sequence replay. However, the replay MI is relatively robust to connection loss for up to 75% connection loss (p < 0.001, rank sum test).

As shown in Fig 5c, we plot and compare the stimulation-induced sequence MI for the trained network and untrained network under different background noise levels. For each replay event, a 50 ms time window is applied to calculate the MI value. Note that, the MIs of the sequence replay in the trained PFC network differ significantly from their counterparts in the untrained PFC network at all the noise levels (p value < 0.001, rank sum test). In the trained network, the sequence MI reaches maximum at intermediate noise level. Weak and strong noise levels will both degrade the MI in different ways. In the case of weak noisy input, the membrane potentials of the PY cells are far below the voltage threshold. The PY cells are less likely to fire, which in turn decreases the number of PY cells recruited during the replay. On the other hand, under the strong noisy input, the membrane potentials of PY cells are driven closer to the voltage threshold. This effectively increases the randomness of firing for PY cells, which degrades the ordered firing and lower the MI value. Therefore, the intermediate background noise level balances the tradeoff between sufficiently large membrane potentials and the randomness of firing, which leads to the most ordered replay and the highest MI value. In the untrained network, for the low noise level cases, the firings of the first 10 PY cells are not sufficient to induce firing in the downstream PY cells, which leads to MI value of 0. However, as the noise level increases, the membrane potentials of the downstream PY cells are depolarized so that the firing of the first 10 PY cells will drive part of the PY cells to fire. Therefore, the MI starts to increase. These results indicate that a proper spontaneous background noise level will affect the sequence replay and is very important in the memory retrieval process.

In order to examine the effect of AMPA connection loss on the sequence replay, we eliminate a random number of AMPA connections in both the trained and untrained PFC network by setting the conductance to 0 and compute the MI value under the same stimulation protocol as before. The resulting sequence MIs under different proportion of AMPA connection loss is shown in Fig 5d. In the trained network, as more and more AMPA connections are eliminated, the MI value gets lower because of less PY cell firings due to the decreasing number of EPSPs from the presynaptic PY cells. On the other hand, the sequence MI is actually robust to recurrent AMPA connection loss. Even though the MI value of sequence replay drops with an increasing loss of AMPA connections, the MI value for the trained network is still significantly different from that of the untrained network for up to 75 % AMPA connection loss (p-value < 0.001, rank sum test). These results indicate that under abundant recurrent connections, the trained network is relatively robust to connectivity damage or degradation in order to successfully recall the memory.

### The sequence replay in PFC network induced by non-specific background noise

Besides the stimulus-induced sequence replay discussed above, we also investigate the spontaneous replay induced by non-specific background noises to all the PY cells in both the trained and untrained network. In other words, we want to examine the possibility of emerging memory retrieval without specific external cues. The non-specific spontaneous noise induced replay resembles the experimentally observed spontaneous reactivation events in the neocortex, where no specific inputs are present [21-23]. The generation and control of these spontaneously emerging replays could be attributed to the neuromodulator system, which exerts a widespread modulation of the overall excitability in a large neuronal network [24, 25].

To simulate this, we adjust the background noise parameters to the PY cells. In this case, the PY cells in the PFC network do not receive any cell-specific stimulation. Each PY cell in the PFC network receives statistically the same background noise. The raster plot of the trained network from one representative simulation (noise level = 0.052 nA^2^) is shown in Fig 6a. The raster plot of the untrained network under the same simulation condition is shown in Fig 6b. It can be seen that under the non-specific spontaneous noisy input, the sequence replay can indeed emerge randomly for the trained network. However, compared with the sequence replay induced by cell-specific current injection, the sequence length and the initiating neurons for each recall can vary. For the untrained network, the noise induced sequence replay is not possible. Similar with the cell-specific stimulation case, we quantitatively investigate two parameters that might affect the MI under spontaneous noisy input, namely the AMPA noisy input level and the AMPA connection loss.

**Figure 6.**
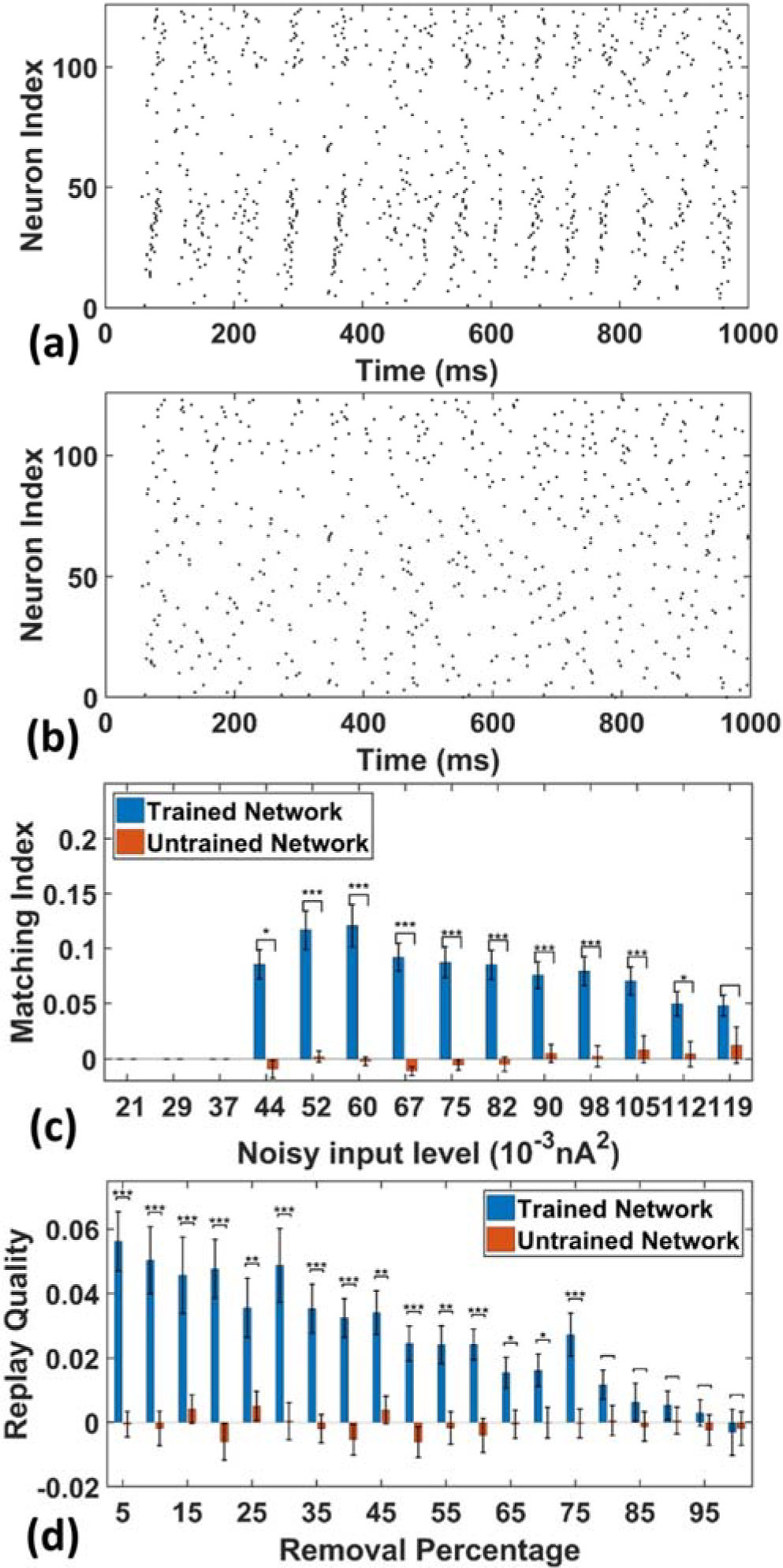
The sequence replay in PFC network induced by non-specific spontaneous noisy input. (a) Under the noisy input to the trained PFC network, the sequence emerges randomly among the PY cell populations with a varying replay length and initiating neurons (noise level = 0.052 nA^2^). (b) For the untrained PFC network, the firing activity is random and no sequence replay emerges. (c) The MIs of sequence replay in the trained network for different noisy input levels. For low noisy inputs, sequence replay cannot emerge. The MI value reaches the maximum for intermediate noisy input levels. For high noise level, the sequence replay in the trained network do not differ significantly from that in the untrained one. (d) The MIs of sequence replay in the trained network for different AMPA synapse losses. Even though the MIs are low, they still differ significantly from those in the untrained PFC network.

As shown in Fig 6c, the sequence MIs are compared between the trained and untrained PFC network for different noisy input levels. For the trained PFC network at low noisy input level, the MI is very small. As the noise level increases, sequence replay starts to emerge and the MI reaches the maximum value and then drops for larger noise levels. This is because when the noise level is low, the membrane potential of the PY cells are far below the voltage thresholds. Therefore, the firing of PY cells is relatively sparse and an ordered replay that recruits a large number of PY cells is hard to achieve, leading to a low MI. When the noise increases to an optimal intermediate value, the membrane potential is elevated to a sufficient level and the firing of certain PY cells is enough to induce the firing of downstream PY cells. If the noise further increase to an overly high level, the randomness firing overwhelms the network and the MI degrades. For the untrained network, the MI values remain low for all the noisy input levels. Note that, even though the MI values for the trained network under spontaneous noisy input are smaller than that for the previously discussed cell-specific input case, the MIs are still significantly different from those for untrained network (p<0.001, rank sum test). These results show that given a proper noise level, the memory replay is indeed possible under non-specific background noise in the trained PFC network.

For the AMPA connection degradation condition, similar to the cell-specific case, we randomly delete the same AMPA synapses between PY cells in both the trained and untrained network. Then, spontaneous noisy inputs are given to all the PY cells in the PFC network and sequence MIs are computed. As shown in Fig 6d, for the trained network, the MI drops as the number of removed AMPA connections increases, since the total EPSPs from presynaptic PY cells to one postsynaptic PY cell are decreasing with more AMPA synapse losses. However, for the untrained network, the MI values do not change much and remain low for different AMPA connection losses. Still, the network is relatively robust to connection loss, meaning that the MI values for trained network are significantly different from those in the untrained network for up to 75% AMPA connection loss.

### The effect of SWR number and background noise on memory retrieval

It has been reported that during the early stage of PFC engram cell formation, the natural cues cannot induce the memory recall. However, the engram cells can be activated by optogenetic stimulations, which will lead to successful memory retrieval [26, 27]. Also, the spine density of PFC engram cells in the later days of learning is significantly higher than the early days [26]. These findings lead to the postulation of “silent engrams” and “active engrams”, which refer to the neurons that encode memory, having weak or strong synaptic connections between them respectively. To explore this idea from a computational perspective, we simulate the network for two different numbers of SWRs during training (2000 ms, 10 SWRs and 3800 ms, 19 SWRs) and test the sequence replay using strong current injection to the first 10 PY cells.

The resulting AMPA recurrent connection matrix and the sequence replay are shown in Fig 7. It can be seen that for the shorter training time (2000 ms), the feed-forward connections in the PFC network are already established through LTP (Fig 7a). However, under the stimulation to the first 10 PY cells, the sequence cannot be retrieved because of the weak AMPA connections that are not sufficient to depolarize the membrane potential (Fig 7b). For the longer training time, the AMPA connections among the PY cells in PFC are further potentiated (Fig 7c) and the sequence can be replayed by the same stimulation on the first 10 PY cells (Fig 7d). These results support the idea that the “silent engrams” can get mature with longer training and eventually turn into “active engrams” for successful memory retrieval.

**Figure 7.**
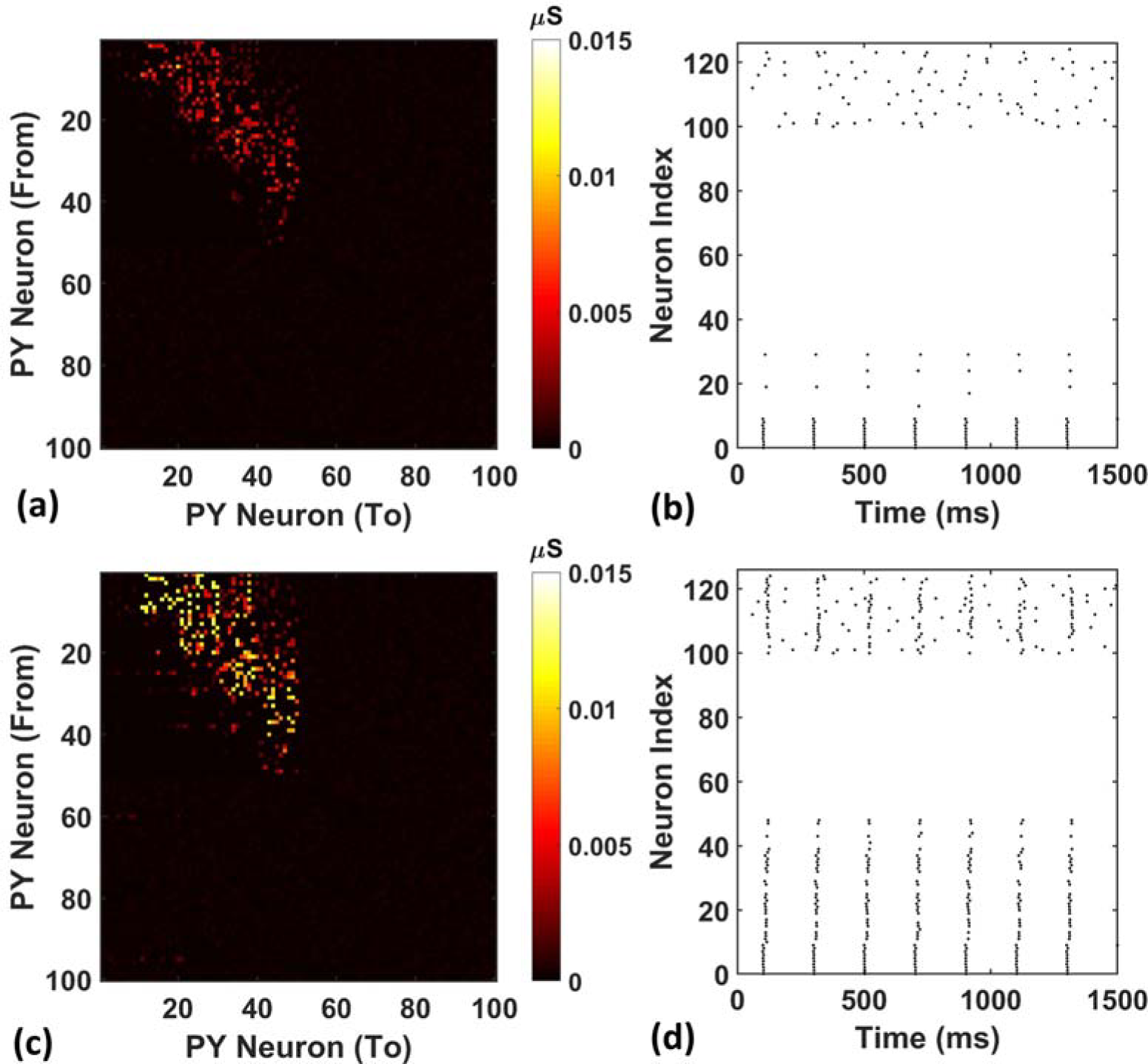
Weight matrices and raster plots for silent engram assembly and active engram assembly under cell-specific strong input. (a) The recurrent AMPA connection matrix between PY cells in the PFC network after short training (2000 ms). The feed-forward connections are strengthened, but not strong. (b) The raster plot of the PFC network under cell specific current injection. The firing of the first 10 PY cells are not sufficient to induce massive firing of the downstream PY cells. (c) The recurrent AMPA connection matrix between PY cells in the PFC network after long training (3800 ms). The feed-forward connections are stronger compared with (a). (d) The raster plot of the PFC network under same input as (b). The firing of the first 10 PY successfully induce a sequence replay in the downstream PY cells.

Further, we investigated the possibility whether we could actually achieve successful memory recall among the “silent engrams”. To simulate this, during memory recall, we apply different background noise levels to the PY cells in the PFC networks going through different training times. The result is shown in Fig 8. The red color indicates successful sequence replay with high MI values, whereas the blue color stands for poor sequence replay with low MI values. It can be seen that indeed, for different noise levels, longer training time will typically result in better sequence replays. Also, by increasing the background noise levels during memory retrieval, good sequence replay can be achieved even for shorter training time. The elevated background noise level effectively increases the average membrane potential and hence the excitability of PY cells. This can be achieved either naturally through neuromodulator system [25, 28] or manually through transcranial electrical stimulation [29, 30]. However, the boosting effect of the noisy inputs does not extend for high noise levels. Further increasing the noise level will hurt the sequence replay and result in low MI values due to excessive firing dominated by randomness.

**Figure 8.**
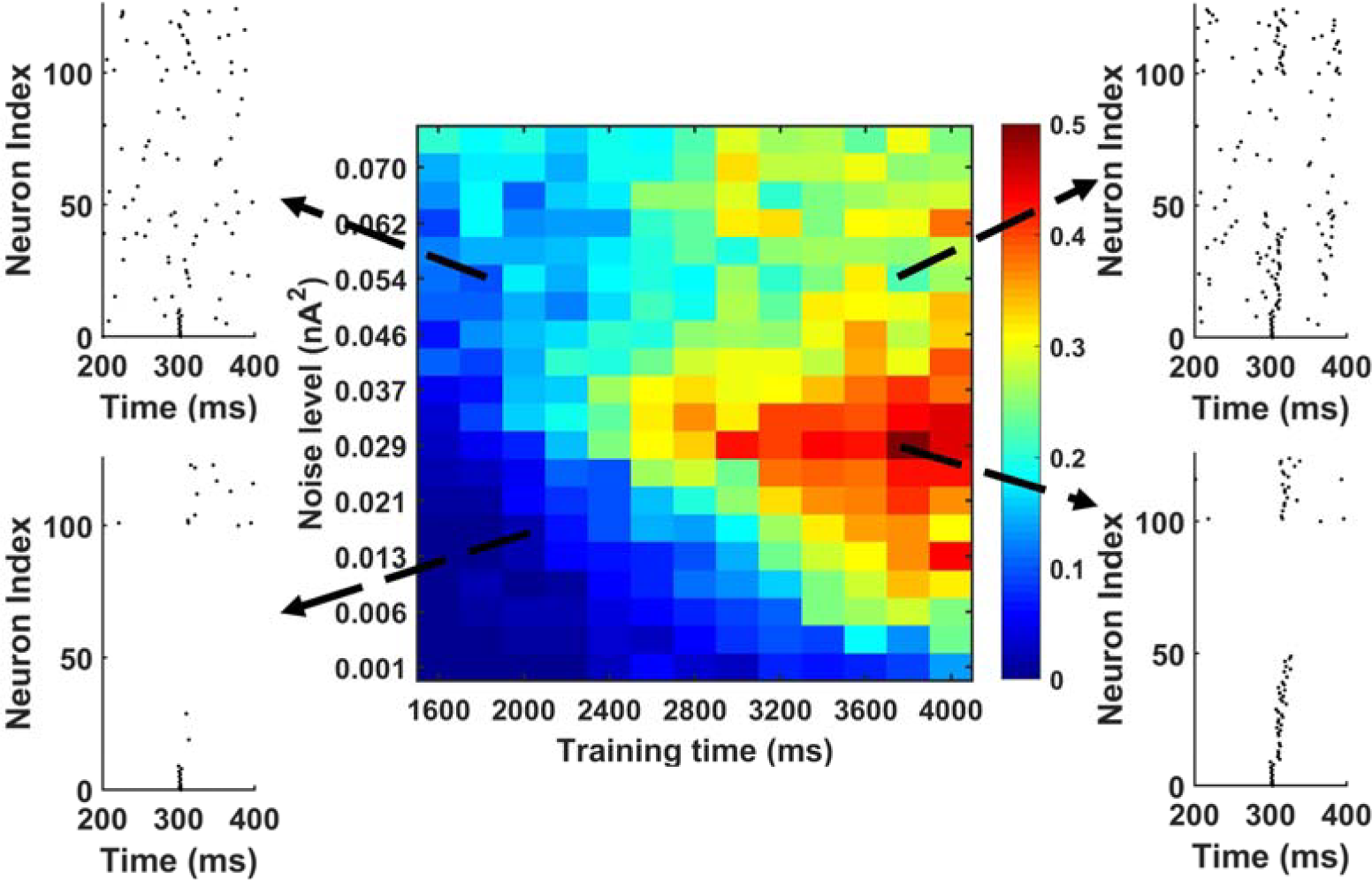
The MI as a function of training time and noise level during memory replay. The first 10 PY cells get strong current injection to initiate the replay. In general, under fixed background noise level, the longer training time is, the better the replay will be. Under fixed training time, moving from low noise to intermediate noise level improves the sequence replay. However, further increasing the noise level will degrade the MI. Also, by increasing the noise level from low to intermediate level, the same sequence MI can be achieved using shorter training time (See the solid black arrow).

## Methods

### Hippocampus model

The hippocampus CA1 model is based on the previous work, which has demonstrated the generation of sharp-wave-ripples under noisy inputs [14-16]. Our adapted model consists of 400 pyramidal cells (PY cells) and 100 basket cells (BS cells), which are chosen to comply with the CA1 neuroanatomy [31]. The pyramidal cell model has five compartments, namely the soma, the basal dendrite, and three apical dendrites. The basket cell is modeled as a three-compartment soma. The geometry of each compartment is listed in Table 1.

**Table 1.**
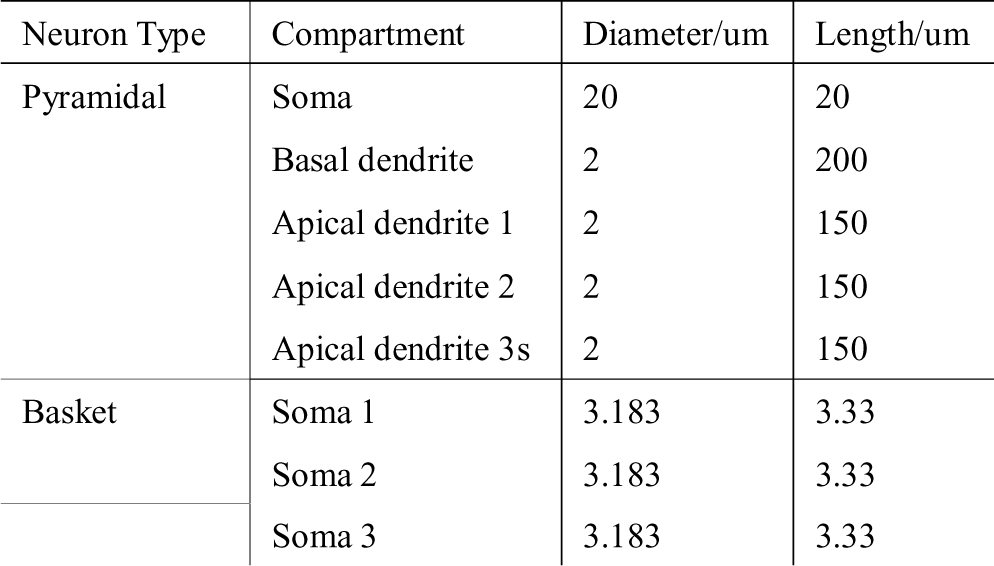
CA1 neuron model.

The connections between different neurons in CA1 network are listed in Table 2. Based on the anatomical study of the connectivity in CA1 area [31], each BS cell receives excitatory AMPA projections from the nearest 50 PY cells, while each PY cell receives inhibitory GABA projections from the nearest 20 BS cells. The BS cell also forms gap junctions with the nearest 4 BS cells. The specifics of the synapse location, strength, and delay are also summarized in Table 2.

**Table 2.**
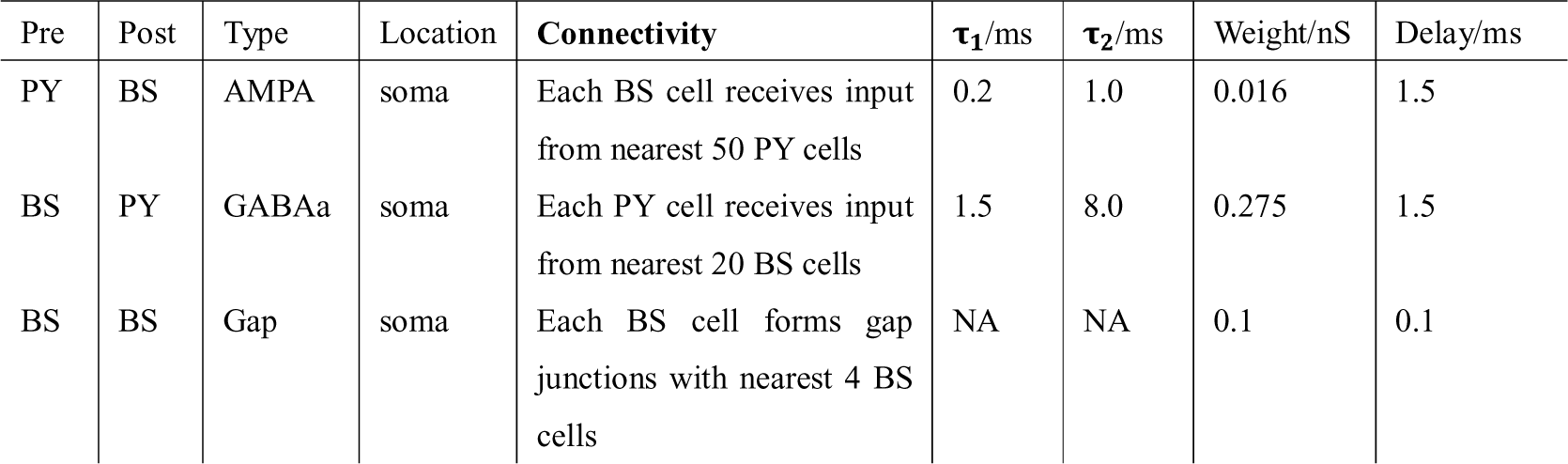
CA1 network connections.

In our CA1 network model, 400 PY cells are randomly allocated into 5 different groups, each of which has 80 PY cells and represents one place cell assembly, as observed in experimental studies [32]. To mimic the synaptic input from CA3, each place cell assembly receives sequential Poisson noisy input. The duration of the noisy input is 35 ms and the start times of two sequential noisy inputs are shifted in time by 15 ms. In this case, each ripple event is driven by the noisy input train that lasts for ~95ms. The setting of this CA3 input pattern is based on recent experimental studies, which show that during awake SWR associated replay, the decoded trajectory from the firing activities of CA1 place cells exhibit discrete “jump” behaviors, which is believed to result from the attractor state changes in the upstream CA3 [33]. Since the mean time for the decoded location change during one ripple event falls within the 25-50 Hz slow gamma range [33], we choose to set the duration of each noisy input to be 35 ms..

### Prefrontal cortex model

Our model for prefrontal cortex (PFC) is based on the layer V PFC microcircuit model published in [17, 34]. We modified the original model to include 100 pyramidal (PY) cells and 25 interneurons (IN). The biophysical properties of the single neuron model are kept the same. Similar to the CA1 network model, the PY cells and IN cells in PFC model also form two 2D planes with 10um spacing and 20 um spacing respectively. The physical specifics of the cell model is shown in Table 3. To reduce the computational complexity of the network and to be consistent with our CA1 model, we reduce the segment number of each compartment to 1.

**Table 3.**
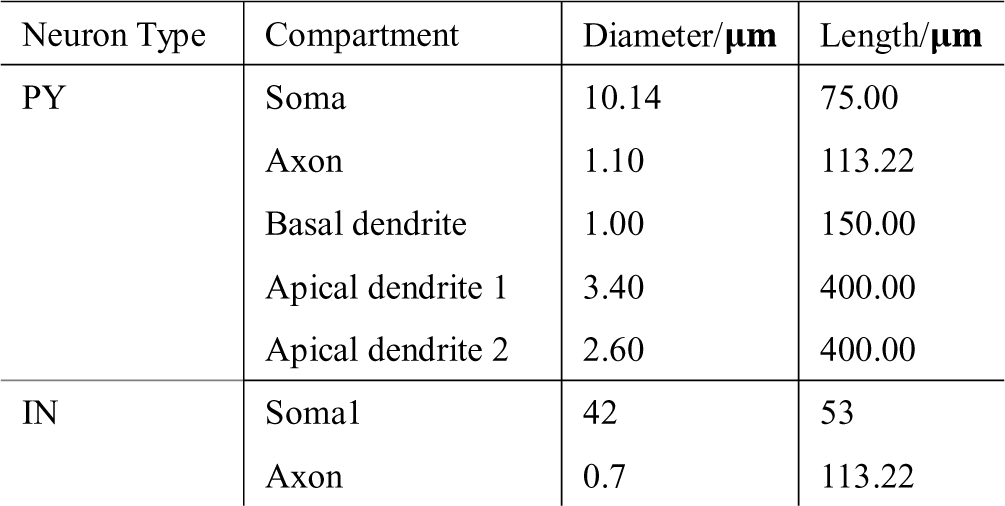
PFC pyramidal cell and interneuron physical properties.

The connections between the cells in the PFC network is shown in Table 4. It has been reported that one PY cell in rat visual cortex makes unidirectional and bidirectional connections to other PY cells with a probability of 0.13 and 0.06 [35]. Since the PFC area has abundant recurrent PY-PY connections, which are more than double the rate than in visual cortex [36], we set the probability of unidirectional connection to 0.25 and the probability of bidirectional connection to 0.12. For the IN cells, each of them is connected to randomly chosen 10 IN cells based on previous anatomical studies [37].

**Table 4.**
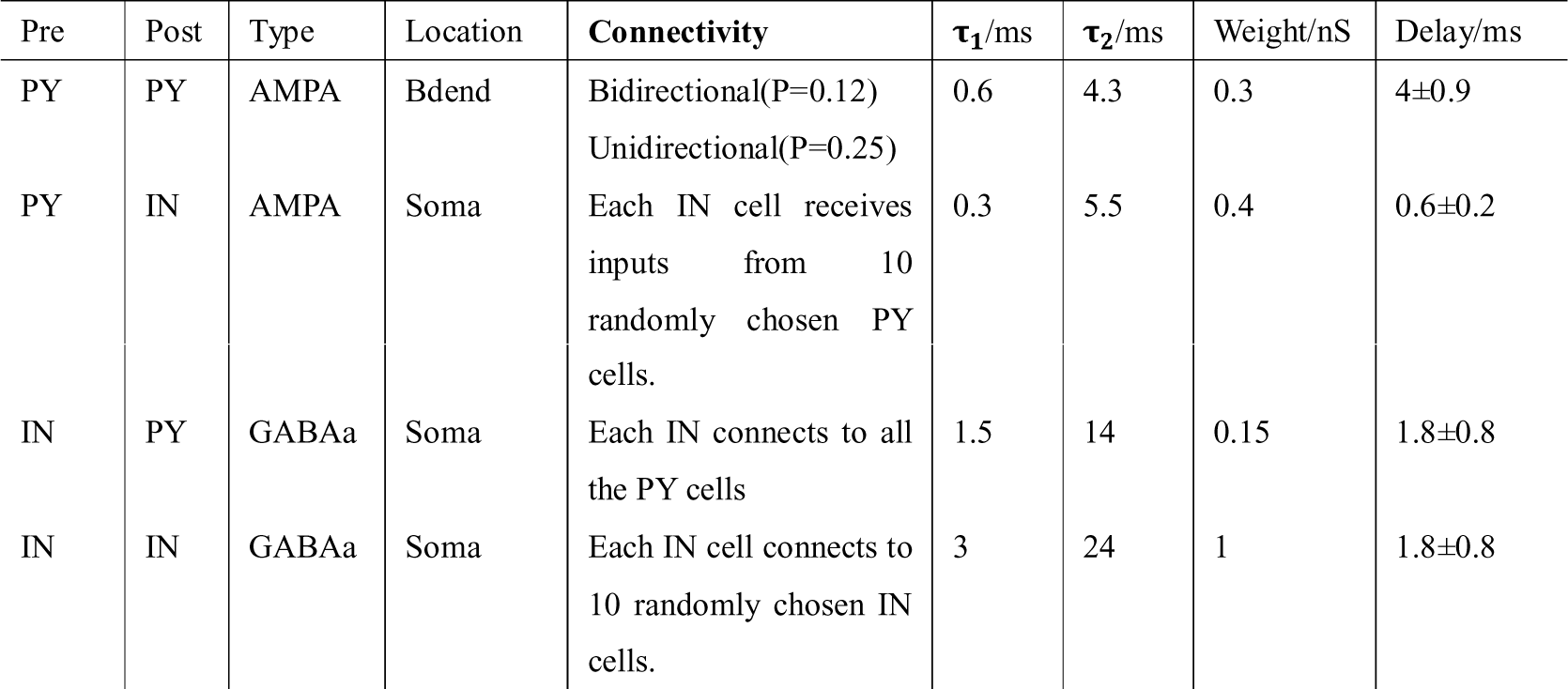
PFC network connections.

During wakefulness, the membrane potentials of the cortical neurons undergo persistent depolarization, which is quite different from that during asleep or anesthetized states [28]. To simulate the background noise on the PY cells and IN cells in PFC, we add independent Poisson inputs to the dendrites of PY cells and soma of IN cells. As a result, the PY cells and IN cells exhibit stochastic firing activities with frequencies of 0.11 Hz and 9.25 Hz respectively, which are consistent with the reported PFC cells firing rates [38, 39].

### CA1 and PFC connectivity

It is well known that the hippocampus closely interacts with a large number of cortex areas, including the PFC, in both monosynaptic and multisynaptic pathways [18, 40-42]. The ventral hippocampus CA1 region and the proximal subiculum make monosynaptic projections directly to both the excitatory and inhibitory neurons in PFC [43-45]. Based on these experimental findings, the pyramidal cells in our CA1 model project to both the pyramidal cells and interneurons in PFC via AMPA synapses. The specifics of the connection are shown in Table 5. In the PFC model, 50 pyramidal cells are randomly chosen and divided into 5 different groups. Each PFC PY cell in one group receives AMPA projections from randomly selected 30 out of 80 PY cells from a specific place cell assembly in CA1. Similarly, the selected CA1 PY cells also form AMPA synapses to all the IN cells in PFC for feed-forward inhibition.

**Table 5.**
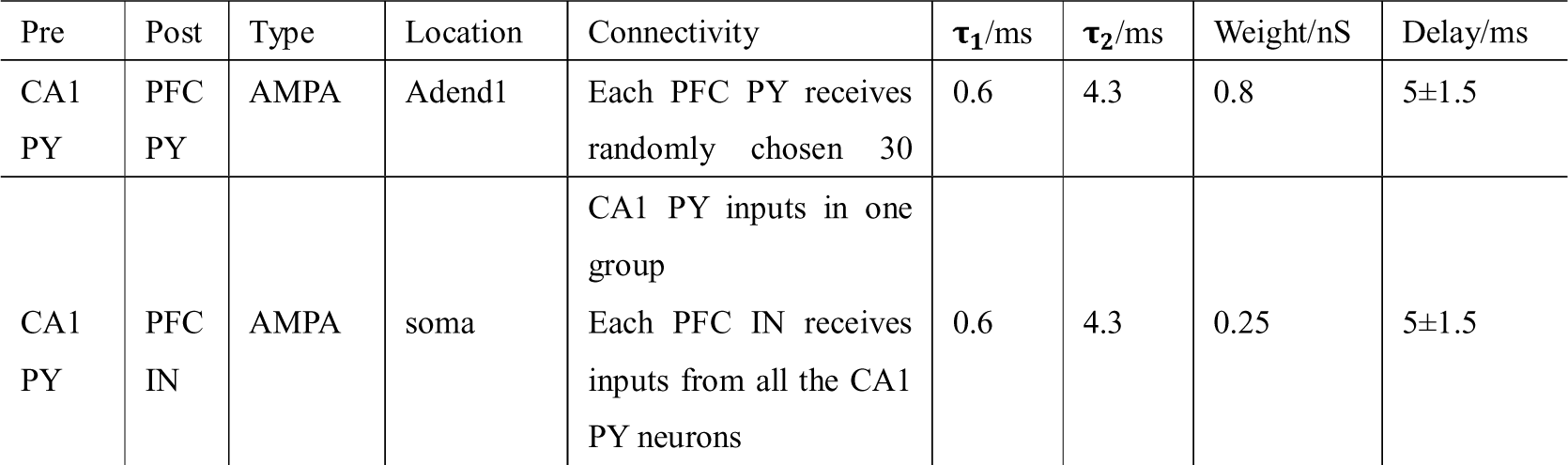
CA1-PFC interconnections.

### Calculation of background noise

In our model, the background noise of each neuron is introduced through the noisy AMPA synapses of Poissonian characteristics. In order to quantify the intensity of the background noise, we compute the second moment of the noise current, which is the same as previous studies [14, 16].

### Spiking-time dependent plasticity

To investigate the sequence learning capability of the cortex model under CA1 input, we implement STDP for both the PY-PY AMPA synapses and IN-PY GABAa synapses to make sure that the excitation-inhibition balance is maintained throughout the learning process [46]. The excitatory STDP for PY-PY AMPA synapses has a classic asymmetric shape [47], while the inhibitory STDP for GABAa synapses has a symmetric shape, which has been reported recently in layer V cortical network [48]. The formula for the STDP of PY-PY AMPA synapses is shown in (1), *W* where is the current AMPA synaptic strength; is *W*_*TH*1_ the target LTP synaptic strength for AMPA synapses; *p*_1_ is the potentiation factor; *τ*_*p*1_ is the LTP time constant; *τ_d_* is the LTD time constant. The formula for the STDP of IN-PY GABAa synapses is shown in (2), where *W* is the current GABAa synaptic strength; *W*_*TH*2_ is the target LTP synaptic strength for GABAa synapses; *p*_2_ is the potentiation factor; *τ*_*p*2_ is the LTP time constant. To prevent divergence, the synaptic weights for AMPA and GABAa synapses are restricted in a defined range shown in (1) and (2).

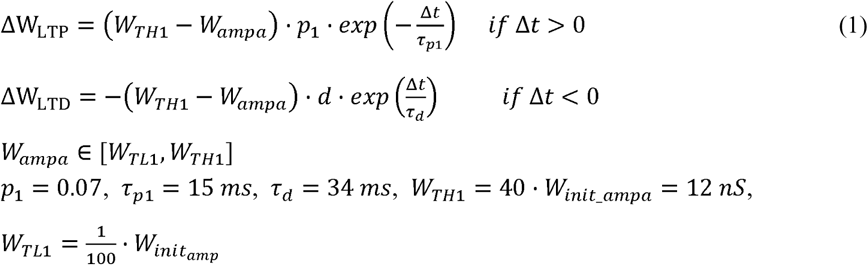

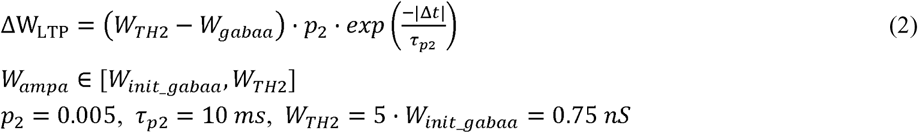

### Sequence replay analysis

To quantify the sequence replay, similar with previous studies [3, 49], we define a matching index (MI) shown in (3). In this equation, N is the total number of neuron pairs from different groups. Since there are five groups in the simulation, each of which have 10 neurons, the value of N equals 1000. n_correct_ is the number of correctly ordered neuron pairs between any two groups and n_wrong_ is the reversely ordered neuron pairs. When all the neurons fire and form a sequence in the ideal order (group A -> group B -> group C -> group D -> group E), the term n_correct_ will equal to N and n_wrong_ equals to 0. Therefore, the value MI will be 1. For ideal reverse replay in the form (group E -> group D -> group C -> group B -> group A), the term n_correct_ will equal to 0 and n_wrong_ equals to N. Therefore, the value MI will be −1. If the sequence replay is totally random, on average, the term n_correct_ and n_wrong_ will be equal. In this case, the value MI will be close to 0.

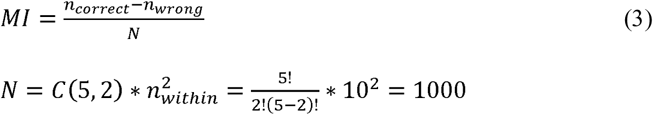

### Calculation of local field potential

To compute the local field potential, we sum up the total electric field generated by the transmembrane and postsynaptic currents across all the compartments of the neurons[50]. The recorded electric potential is computed by equation (4) using the source-field model of current monopoles. In the equation, ϕ(r, t). is the recorded potential at time t and position r. σ is the extracellular conductivity. N is the total number of compartments. |r – r_n_| is the distance between the compartment n and the recording position.

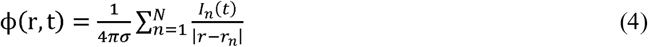

## Acknowledgments

We would like to thank Yuhan Shi for useful discussions.

## Data Availability Statement

The model is available from the ModelDB database (http://xxxx) with accession number xxx. All other relevant data are provided in the manuscript.

